# Neural network-assisted analysis of tube vocal tract models

**DOI:** 10.1101/2025.08.19.671092

**Authors:** Runhui Song, Johan Sjons, Axel Ekström

## Abstract

We present a pipeline for deep neural network assisted modeling and analysis of tube vocal tract models. Such models are composed of a series of cylindrical tube segments, each characterized by length and cross-sectional area. A large synthetic dataset of such tube configurations is generated, and a circuit theory–based algorithm predicts corresponding formant frequencies. To explore the mapping between tube sequence shapes and predicted resonance (formant) values, the pipeline integrates both linear regression and nonlinear machine learning models —including multi-layer perceptrons. Model interpretability is assessed using Shapley Additive Explanations (SHAP), which quantifies the contribution of each segment to predicted formant frequencies. The proposed framework enables detailed exploration of the articulatory-acoustic relationships inherent to an acoustic tube and vocal tract simulacrum. We present and describe the pipeline in the context of modeling effects of perturbations on the first three predicted resonances for a 16- cm tube, divided into 1 cm segments. Our pipeline can be applied to any method that models predictions of behavior of an acoustic tube, where the tube is conceived as a series of segmented units.

## Introduction

In any acoustic resonator, its shape and dimensions determine specific resonant frequencies at which sound waves reinforce one another. This physical process is the basis for various phenomena in speech production, where spectral energy peaks are termed formants (1). Essentially, the vocal tract functions as a resonator that dynamically changes shape with the movements of various articulators (lips, tongue tip, tongue body, etc.) (2–4). For this reason, the vocal tract can be represented as an elongated tube – a series of segments defined by their length and cross-sectional area. The idea as such is long and well established in phonetic sciences (3, 5, 6), with modern implementations going back some 75 years (2, 7). Today, computational implementations of tube acoustics make possible the investigation of complex theoretical articulatory-acoustic phenomena (5, 6, 8). We present a pipeline, DeepVocalTube (DVT), for the analysis of large such datasets using deep neural networks (DNNs).^1^

It is long established in phonetic sciences that the first three resonant frequencies (hereafter formants, *F*_1_ *F*_2_ *F*_3_) correspond well to vowel quality (3, 5, 6). As such, the single-most crucial sanity test for any large-scale modeling attempt of tube acoustics is the accurate modeling of these three formants. Below, we describe a simplistic use case for our pipeline – exploring the relative contributions of each segment in a sequence, to predicted first, second, and third formants.

### Simulating the behavior of an acoustic tube

In theory, several different algorithms may be used to derive predicted formant frequencies (4, 9–11). The purpose of our pipeline is not empirical investigation *per se*, but methodological innovation. In theory, our pipeline applies to any vocal tract modeling effort, where a model is constructed of segments defined by length and area. As such, in this work, we opted for the approach developed by Fant, for simulating the behavior of a lossless closed-to-open tube (3, 4, 8). The algorithm is both simplistic (making few assumptions), computationally efficient, and well established in the literature (3). Specifically, we here use the computer program presented by Liljencrants and Fant (1975), which simulates digitally behavior observed in analogue circuits (2, 3, 7, 12), with predicted formant frequencies calculated based on a transfer function from glottis to lips.

The algorithm^2^ recursively computes a determinant through tube segments, the value of which is called the *transfer determinant* Δ_*n*_, reflecting the impedance transformation up to the *n*_*th*_ tube segment. The angular frequency *ω*, measured in radians per second (*rad/s*) is defined as:

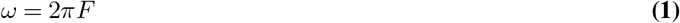

where *F* represents the frequency of the sound wave measured in Hertz (*Hz*), which corresponds to the specific acoustic frequency being simulated. For example, if the response of the tube to a 500 *Hz* sound wave is studied, then *F* = 500 *Hz*. The normalized phase angle of the *n*_*th*_ tube segment is:

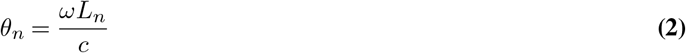

where *c* = 35300 cm/s is the speed of sound at 35*°C*, and *L*_*n*_ is the length of the *n*_*th*_ segment. The ratio of the area of two connected tube segments (*A*_*n*+1_ *and A*_*n*_) is represented as:

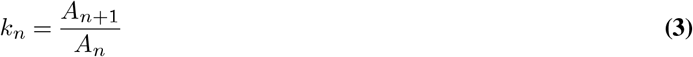

The recursive formula for the transfer determinant is:

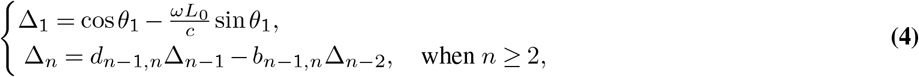

where:

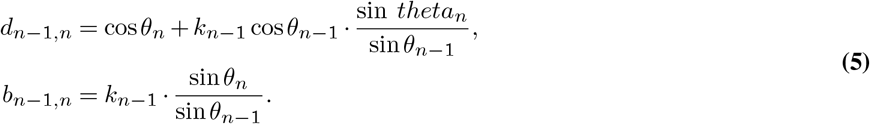

After obtaining the determinant of the final tube segment Δ_*M*_, a quasi-spectral function is constructed:

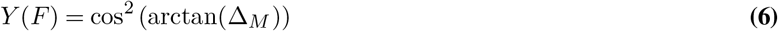

DVT also includes an internal end correction for segments where a narrow segment opens into a much wider segment. The original computational approach developed by Liljencrants and Fant (1975, p. 16) did not include such a correction, “since actual [vocal tract] configurations seldom display such discontinuities”. However, because DVT models the behavior of an acoustic tube – including but *not* limited to realistic vocal tract configurations – we determined such an end correction was necessary. The correction was implemented based on Ingard (1953) and Fant (1971). It determines that where the relationship between a smaller segment *A*_0_ and a more expansive segment *A* is *A*_0_ *<* 0.16*A*, the length of *A*_0_ may be adjusted to compensate. This change is expressed as:

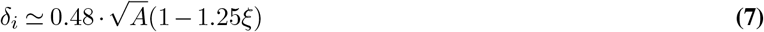

where *A* denotes the cross-sectional area of the greater-area section, and 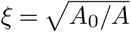 is a correction factor indicating the ratio of curvature to area. Note that depending on the purposes of modeling, additional internal end corrections (14, 15) may be similarly included in data generation procedures.

### Dataset Generation

It is generally agreed upon that while a small number of tube segments allow for simplistic modeling of influential factors such as opening (jaw and lips), tongue position, and pharyngeal constriction into account (3), additional segments allow for finer discrimination and more accurate representation of vocal tract behavior (3, 6, 7, 16). For example, the “distinctive regions model” developed by Mrayati and colleagues (1988) identifies eight regions necessary to capture naturalistic speech-like behavior of a tube (where each region corresponds to a selectively “sensitive” part of the tract, where changes are disruptive and selectively influential on formant output). In addition, the reliability of DNNs is contingent on achieving close fit to data, even when data becomes more complex. For these reasons – to showcase both the appropriateness of the modeling procedure to basic research in speech production, and the applicability of our pipeline as benefiting future such research – we constructed a 16-segment model divided into equal-length segments of 1 cm in length (Fig. 1).^3^

**Fig. 1.**
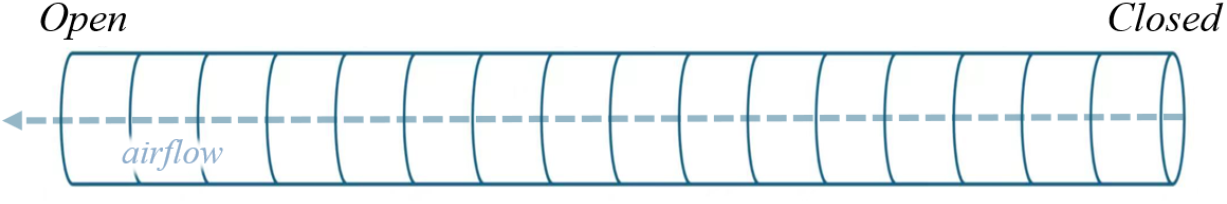
An open-to-closed acoustic tube, segmented into 16 equal-length segments. For comparisons with speech production, the initial “closed” section corresponds to the laryngeal opening, while the “open” segment corresponds to the termination of the oral tract, i.e., the lips.

**Fig. 2.**
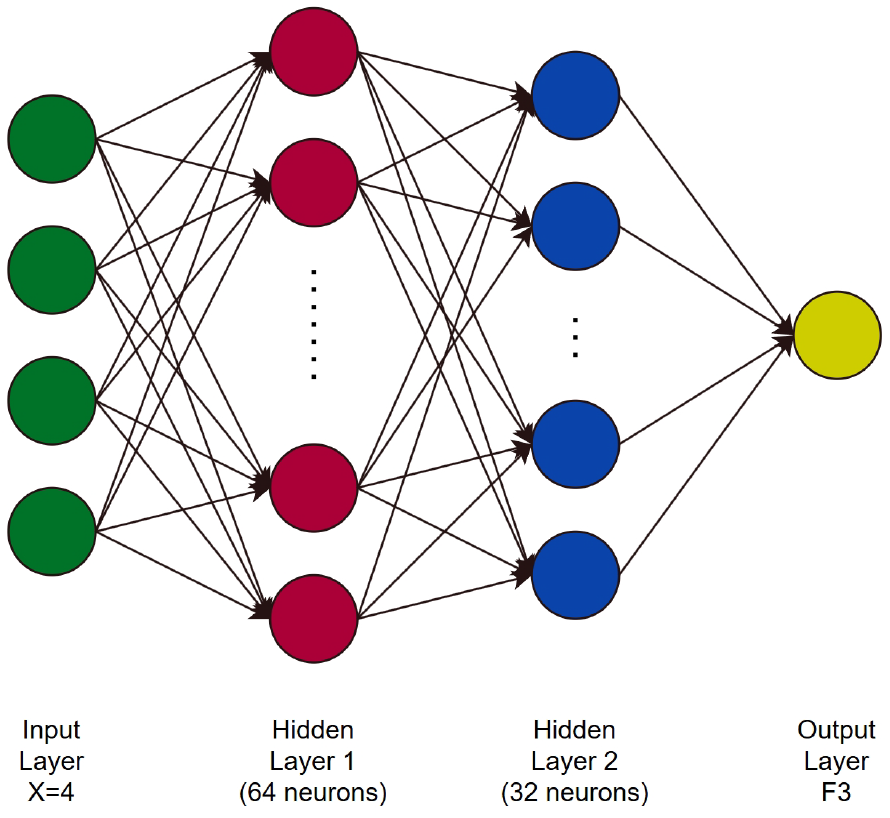
Example illustration of Multi-Layer Perceptron (MLP) structure, with 4 input features.

For training and evaluating the multi-layer perceptron (MLP), we generated 40,000 sample data, corresponding to random permutations of the 16-segment tube sequence, where each segment was varied between 0.1 cm^2^ and 10 cm^2^. To validate our choice of 40,000 samples, we conducted a sensitivity analysis of dataset size by using different amount of data (1,000, 5,000, 10,000, 20,000, 40,000, and 80,000). A larger sample is possible (in theory, up to the total number of possible permutations), provided no computational limitations exist. However, data samples cover a wide range of tube model configurations, enabling the models to learn the relationship between tube shape and formant frequencies. Since the length of each tube segment was kept constant in the modeling, only the area of tube segments was used as an input feature in the subsequent modeling.^4^

## Data Analysis Method

### Linear Regression as Baseline Model

A multiple linear regression was constructed using the nth formant frequency as prediction target and the cross-sectional area of tube segments as the input feature. This model was used as a baseline to assess whether a linear relationship exists between tube configuration and acoustic prediction. However, the model was poorly fitted to the dataset, with *R*^2^ lower than 0.32 across all three formants and residual analysis showing a significant nonlinear relationship between input features and predicted formants. While attempts can be made to transform the inputs or outputs as a function to improve the linear fit, this approach often lacks physical interpretability and may introduce potential bias (18). For this reason, we applied more advanced nonlinear modeling approaches to capture the complex relationship between tube segments and predicted resonance frequencies.

### Machine Learning Approaches

Considering the evidently limited ability of linear models, we used a commonly used non-linear modeling method, the multi-layer perceptron (MLP). This machine learning model is able to effectively capture complex nonlinear relationships, and, given subsequent analysis, provide interpretability. Here, we applied eXtreme Gradient Boost (XGBoost) as a baseline model (19, 20) This method is an integrated learning algorithm based on decision trees, where each new tree is used to correct the residuals of the previous set of models, thus gradually approximating the target output. Both models have been tested with the same data configuration, where MLP proved to have better performances in the 16-segment tube experiment (17).

The structure of our MLP network was set as (1) an input layer containing 16 elements (cross-sectional areas), (2) two hidden layers containing 64 and 32 neurons respectively; (3) an activation function *ReLU*, applied in each hidden layer to introduce nonlinearity; (4) an output layer, a neuron for predicting the continuous variable *N* corresponding to *N* ∈ {1, 2, 3} formants. The MLP was trained using the Adam optimizer (21) with a learning rate of 0.001. Because it represented a regression task, the loss function was the mean squared error (MSE), and the maximum number of training epoches was set as 2000.

### K-Fold Cross-Validation

In order to increase the reliability of the evaluation of the model’s performance, we employed a *k*-fold cross validation. By dividing the dataset into training and test set, it could help to avoid overfitting (22). Specifically, the data were divided equally into *k* non-overlapping subsets (folds); in each iteration, the *k* − 1 folds were used for training, and the remaining one for testing, looping *k* times such that each subset was used as a test set once. The final model performance was then averaged over all folds of evaluation metrics. Here, a 5-fold cross-validation was used, such that four fifths (80%) of all generated data were used for training and, and the remaining fifth (20%) for testing. The reason for using five folds was to increase the model’s ability to generalize (23), while still retaining a sufficiently large training set in each split.

### SHAP for Model Interpretability

Neural models are “black boxes”, precluding insights into how predictions are made (e.g., 24). To facilitate interpretability of the model output, the DVT pipeline employs the Shapley Additive Explanation (SHAP) framework – a method derived from game theory (26) – to uniformly measure the contribution of each input feature to the model prediction results. SHAP measures the average contribution of a given input feature to the overall prediction across all possible sequences of input features.

Specifically, a positive SHAP value indicates that the feature contributes to increasing the predicted value compared to the model’s average output, and a negative SHAP value indicates that the feature contributes to decreasing the prediction. Here, we employed two types of SHAP visualizations, (1) the Summary Plot, which aggregates SHAP values across all samples and features, highlighting feature importance, as well as to what extent high and low values of each feature affect the prediction; and (2) the Waterfall Plot, which illustrates a single prediction by showing how each feature changes the output from a baseline value to the final prediction. In sum, (1) allows for visual inspection of the contributions of every segment to formant output, as observed in the dataset, while (2) visualizes the contribution of segments to formant changes within a single tube model. To facilitate interpretation of the MLP, we adopted KernelExplainer (25) within SHAP, which approximates the marginal contributions of features by perturbing model inputs and constructing a local linear model. SHAP provides both local and global interpretability, as well as uniform and robust interpretation results for different types of models, which is particularly suitable for the needs of multi-model comparison and structural analysis (27).

### Equipment

Experiments were conducted on a laptop equipped with an AMD Ryzen 7 8845H and 32 GB RAM.

## Evaluation Metrics

We employed a total of five common regression metrics to evaluate model performance: the coefficient of determination (*R*^2^), mean absolute error (MAE), mean absolute percentage error (MAPE), mean square error (MSE), and root mean square error (RMSE). Each metric offers complementary data pertaining to model performance, and together provide a coherent summary of model quality. MAE, MAPE, MSE, and RMSE indicate predictions’ average offset from the data. *R*^2^ indicates how well the model explains variation in the data.

### Coefficient of Determination

*R*^2^ is an estimate of how much of the variance in the target variable is explained by the model. It is defined as:

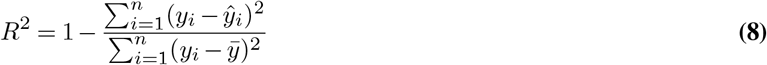

where *y*_*i*_ corresponds to true values, *ŷ*_*i*_ to predicted values, 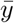 to the mean of *y*_*i*_, and *n* to the number of data points. An *R*^2^ value ranges from 0 to 1, where 1 means perfect prediction, and 0 means the model cannot explain any of the variance in the data (only predict the average). For our purposes, a higher *R*^2^ suggests that a model better captures the relationship between tube configurations and formant frequencies.

### Mean Absolute Error

MAE is the mean absolute difference between the predicted values and the actual values in a dataset. It is calculated as:

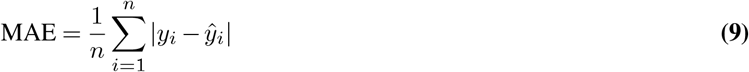

The lower the MAE, the better a model fits a dataset. A MAE value of 0 means the model predicts perfectly. MAE treats all errors equally, without giving extra weight to larger ones, so it provides an average measurement of the model’s performance.

### Mean Absolute Percentage Error

MAPE is the computed average of the absolute relative errors between the predicted and true values, expressed as a percentage:

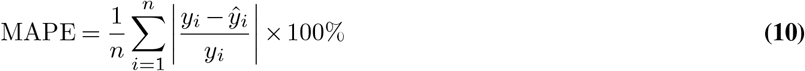

The smaller the value of MAPE, the more accurate a prediction is. For example, a MAPE value of 11.5% means an average difference between predicted and actual value of 11.5%.

### Mean Square Error

MSE is the average squared difference between the predicted values and the actual values in a dataset. It is given by:

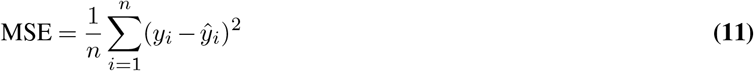

Unlike MAE and MAPE, MSE is more sensitive to observations that are more derived from the mean, thus making it useful for identifying poor predictions.

### Root Mean Square Error

RMSE is the square root of MSE:

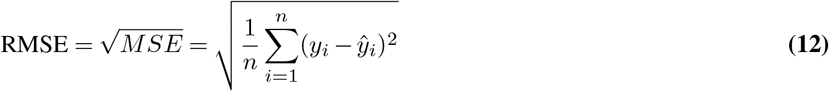

and allows for the observations of model deviations from true labels.

## Results

### Data Sensitivity Analysis

To determine the required data volume for the experiment, we generated eight datasets of increasing sizes: 1,000, 5,000, 10,000, 20,000, 40,000, 80,000, 100,000, and 120,000 samples. We then used these datasets to evaluate the effectiveness of MLP using *F*_3_ – which is more sensitive to minor changes than lower-frequency resonances. The evaluation results for this experiment are presented in Table 1 below.

**Table 1.** Evaluations of model fit for different sizes of dataset.

| Dataset Size | $R^2$ | MAE | MAPE | MSE | RMSE |
| --- | --- | --- | --- | --- | --- |
| 1,000 | 0.3262 | 285.02 | 15.39% | 134478.90 | 366.42 |
| 5,000 | 0.4642 | 251.33 | 13.37% | 108494.82 | 329.30 |
| 10,000 | 0.7072 | 184.48 | 9.88% | 60558.82 | 245.64 |
| 20,000 | 0.8877 | 108.49 | 5.75% | 22757.70 | 150.79 |
| 40,000 | 0.9360 | 78.08 | 4.13% | 13016.64 | 113.96 |
| 80,000 | 0.9521 | 66.27 | 3.51% | 9677.87 | 98.37 |
| 100,000 | 0.9543 | 64.68 | 3.43% | 9316.56 | 96.46 |
| 120,000 | 0.9537 | 64.34 | 3.42% | 9477.86 | 97.27 |

**Table 2.** Evaluations of models fit for each formant.

| Formant | $R^2$ | MAE | MAPE | MSE | RMSE |
| --- | --- | --- | --- | --- | --- |
| F1 | 0.9849 | 8.73 | 2.64% | 144.19 | 12.00 |
| F2 | 0.9768 | 31.37 | 3.02% | 1894.12 | 42.52 |
| F3 | 0.9521 | 66.27 | 3.51% | 9677.87 | 98.37 |

As the dataset size increased, the performance of MLP improved, with *R*^2^ values increasing and all error metrics decreasing. However, the performance gain showed a clear diminishing return: while the increase from 1,000 to 80,000 samples led to large stepwise improvements with each increase, additional benefits of expanding the dataset beyond 80,000 were marginal. For example, the difference in *R*^2^ between 80,000 and 120,000 samples was less than 0.01, and some evaluation metrics results based on 120,000 samples were slightly worse than those based on 100,000 samples. The computational cost also increased sharply: generating 120,000 samples took about 220 seconds compared to only a few seconds for 1,000 samples. The SHAP analysis procedure finished within 1 minute with 1,000 samples, and required nearly an hour for 80,000. Considering both accuracy and efficiency, we chose 80,000 samples for subsequent illustration and investigation, as this size captured the main performance improvements without incurring excessive computational burden.

### Formants

In the following experiment, we investigated how well the MLP model predicts (*F*_1_, *F*_2_, *F*_3_) and how individual tube segments contribute to these predictions. In general, the model achieved high accuracy across all formants, confirming an ability to reliably capture parameter–formant mappings (Tab. 2). To interpret the model, we applied SHAP to analyze the contribution of each tube segment. As an illustrative case, we considered an example tube segment constellation corresponding to a vocal tract configuration for open front unrounded vowel [a], split into 16 segments (Fig. 3).

**Fig. 3.**
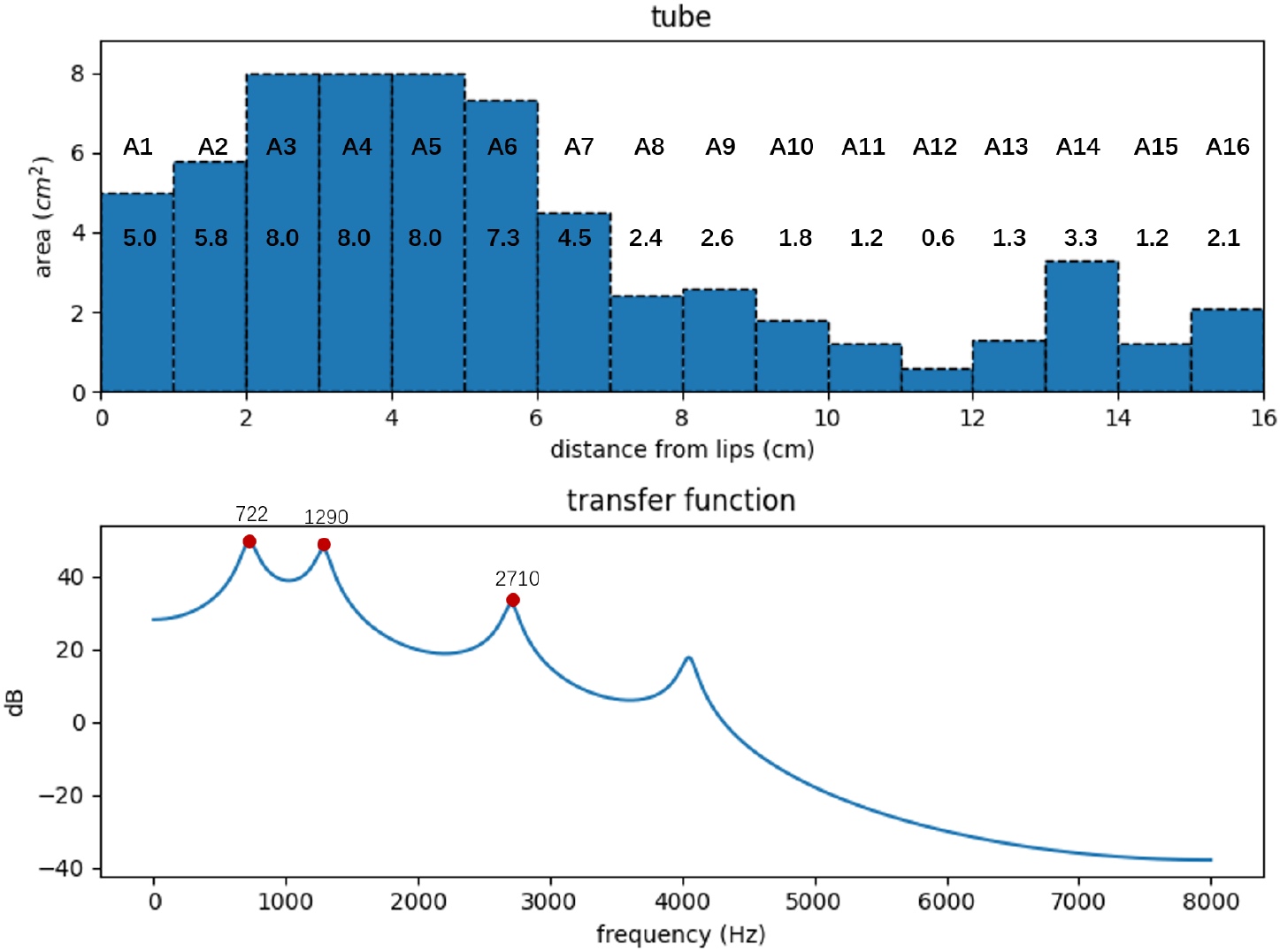
Tube segment parameters and predicted formant frequencies for vowel [a].

Waterfall plots (Fig. 4) illustrate how SHAP explains the prediction for a single configuration. The horizontal axis shows frequency (Hz). The baseline value *E*[*f* (*X*)] represents the dataset average for a given formant (*F*_1_ = 520 Hz, *F*_2_ = 1569 Hz, *F*_3_ = 2638 Hz). Each bar shows the contribution of one segment (*Ai*) relative to this baseline: red bars increase the prediction, blue bars decrease it. The final value *f* (*X*) at the right end represents the MLP prediction for this example ([705, 1300, 2709] Hz), which closely matches the reference prediction from the example ([722, 1290, 2710]). Segments are ordered by contribution magnitude, so the most influential segments appear at the top.

**Fig. 4.**
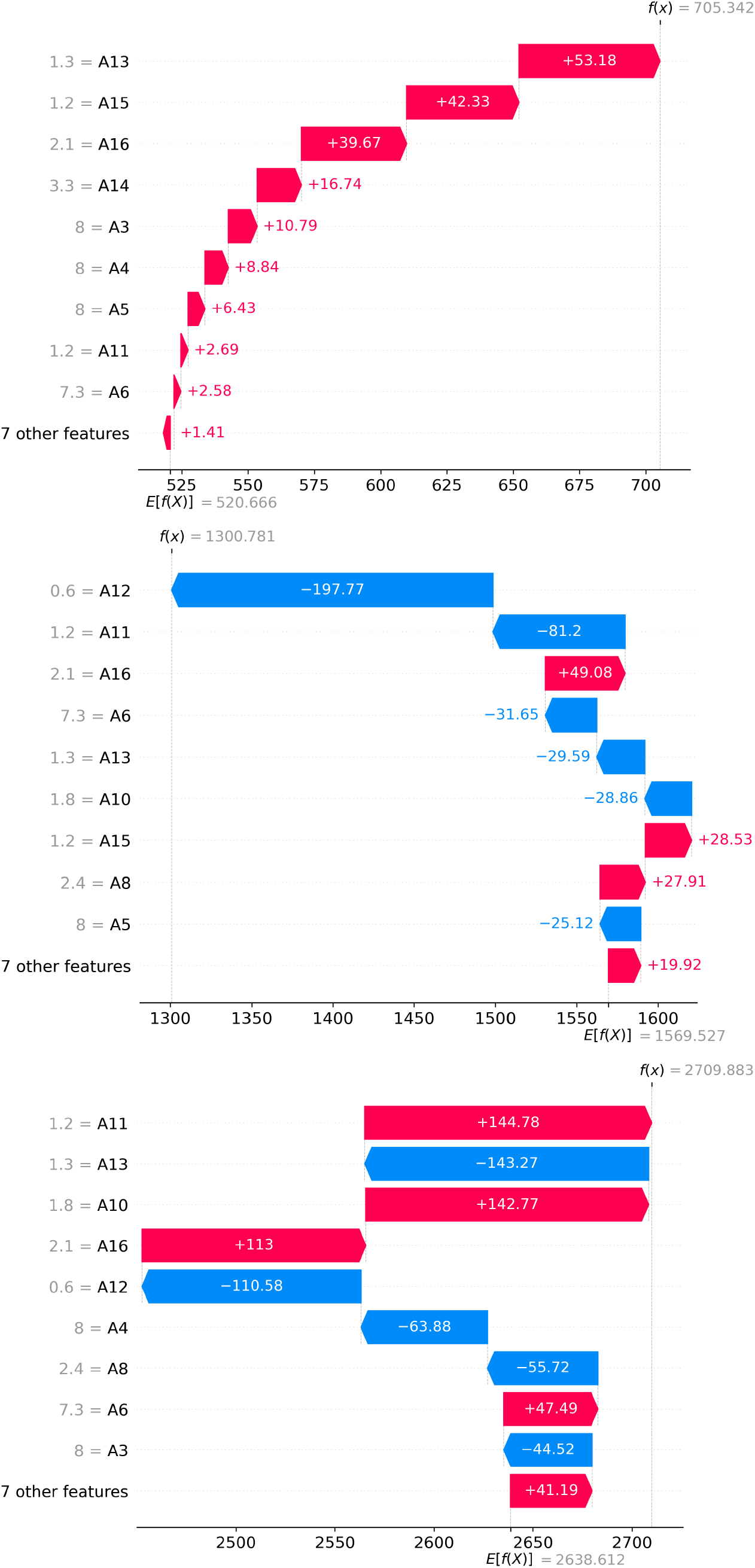
Waterfall plots of SHAP values for predicting F1 (top), F2 (middle), and F3 (bottom) for an example vowel [a], highlighting the contribution of each tube segment in a single example.

The ordering of features in each plot reflects their relative importance. For example, *F*_1_ is strongly influenced by the posterior segments A13, A15, and A16, corresponding to a constricted pharynx raising *F*_1_ in natural speech. *F*_2_ is decreased by constrictions around the mid-back oral cavity A11-A13, while smaller posterior areas A15-A16 increase it. *F*_3_ shows opposing effects between adjacent segments A10-A11 and A12-A13. In contrast to the single-case waterfall plots, the beeswarm summary plots in Figure 5 show the sensitivity of each formant to tube segment variations across the entire dataset.

**Fig. 5.**
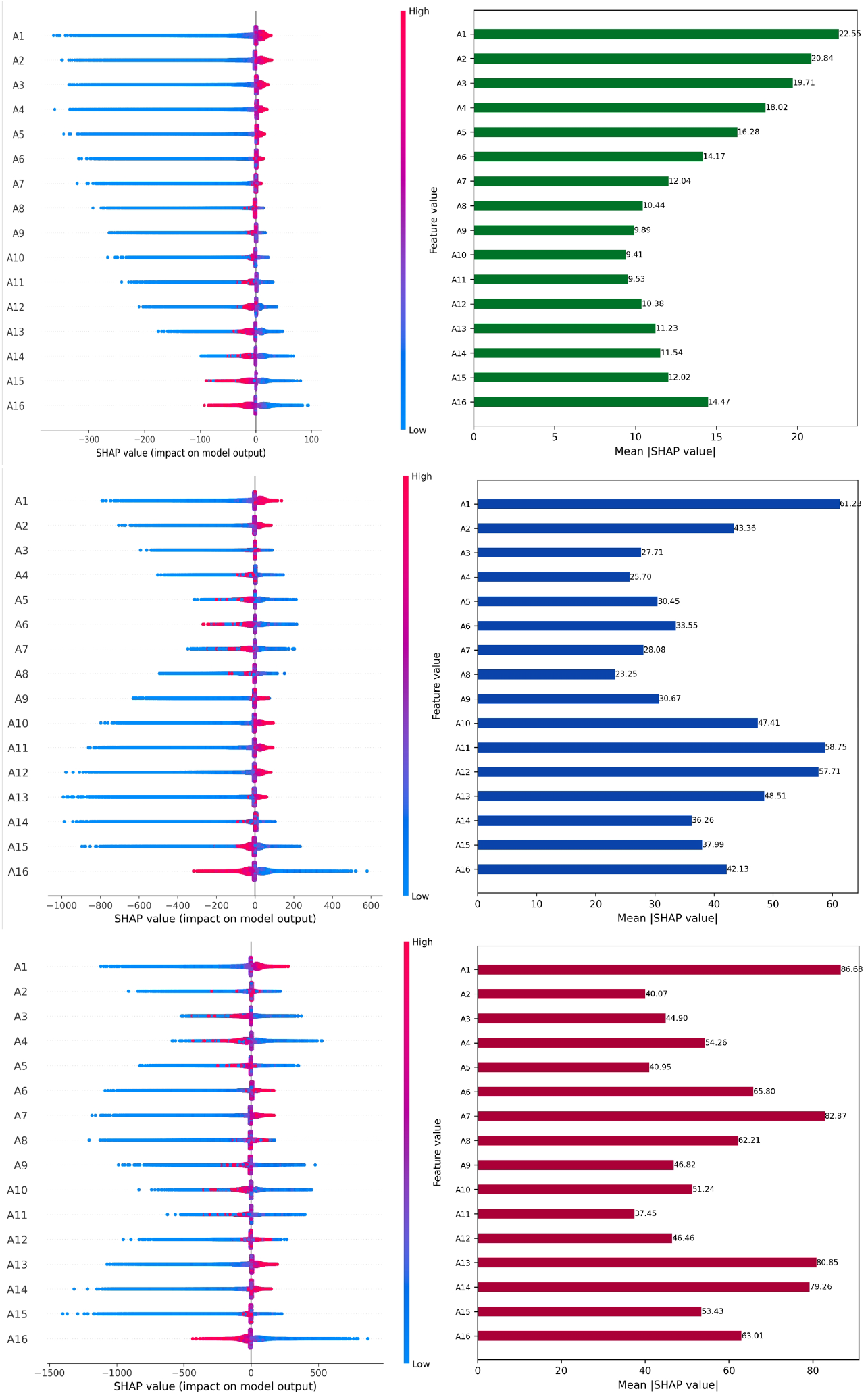
Beeswarm summary plots (left) and bar summary plots (right). The beeswarm summary plots show SHAP values for F1 (top), F2 (middle), and F3 (bottom), illustrating the influence of each tube segment across the entire dataset. Color coding denotes changes to formant frequencies, with blue-colored SHAP values indicating a decrease in segment area, and red-colored SHAP values indicating an increase. Negative SHAP values indicate lower formant frequencies; higher SHAP values indicate higher formant frequencies. The bar summary plots show how large the effect is of each segment by taking the mean of the absolute SHAP values.

For *F*_1_, front sections (corresponding to the lips and the front oral cavity) A1-A5 generally have positive contributions, indicating that widening these regions raises *F*_1_, consistent with earlier studies (3, 5). In contrast, larger posterior areas (corresponding to pharyngeal regions) near the pharynx) A13-A16 tend to lower *F*_1_.

For *F*_2_, effects of segment changes are more complex: Expansion of the anteriormost sections (corresponding to lip opening) A1-A2 raise *F*_2_. Areas corresponding to the oral cavity A3-A8 often show negative SHAP values, where expansion lowers *F*_2_ (28). Expansion in more posterior sections (roughly corresponding to the tongue root area and the front part of the pharynx) A9-A13 again show positive effects, consistent with tongue retraction and “bunching” lowering *F*_2_ in natural speech. Areas corresponding to the lower pharynx A14-A16 have negative effects, such that expansion reduces *F*_2_.

For *F*_3_, several strong effects are observed. The anteriormost section (again, corresponding to the lips) A1 lowers *F*_3_ when constricted, reflecting the typical lip-rounding effect (17, 28). Segments corresponding to the back oral cavity (e.g., A6-A8) have a sustained positive effect on *F*_3_, with A7 being most influential. Segments corresponding to the mid-pharyngeal segments A13-A14 also lower *F*_3_ when constricted. The posteriormost segment (corresponding to glottal region) A16 shows a strong negative effect: when expanded, *F*_3_ decreases.

## Limitations and future work

We have left unexplored interaction effects on formant values between segments. Future modeling endeavors may seek to quantify such nonlinear interactions, for example in the context of verifying or complementing prior theoretical arguments positing distinctively “stable” and “unstable” regions of the vocal tract (6, 16, 29). Such efforts should also be coupled with model selection evaluation (e.g., AIC 30). Given the strive for low computational cost and already high *R*^2^ values, another clear path forward is to include a hyperparameter search for improving the multi-level perceptron model (31).

## Conclusions

The DVT pipeline allows for leveraging tens of thousands of datapoints to answer foundational questions of theoretical phonetic sciences. Our analysis was focused on examining and describing the contributions of individual tube model segments. The software package necessary to replicate and build upon the use cases described here, or to perform any additional experiments, including illustrations, is freely available online.

## Acknowledgements

AE was supported by the Swiss National Science Foundation (PCEFP1_186841, Steven Moran PI). The results of this work and the tools used will be made more widely accessible through the national infrastructure Språkbanken Tal under funding from the Swedish Research Council (2017–00626).

## Footnotes

1 The pipeline and code described in this paper can be accessed through the relevant GitHub repository. https://github.com/folbdsta/DeepVocalTube

2 For full modeling considerations, we refer to the original publications (3, 4).

3 The pipeline in theory applies to any tube model, where predictions are based on parametrized segments (4, 6, 17).

4 Note, however, that the pipeline may be employed to study effects of e.g., elongation of segments.

## Notes

https://github.com/folbdsta/DeepVocalTube

### Competing Interest Statement

The authors have declared no competing interest.

## Bibliography

1. I. R. Titze, R. J. Baken, K. W. Bozeman, S. Granqvist, N. Henrich, C. T. Herbst, and J. Wolfe. Toward a consensus on symbolic notation of harmonics, resonances, and formants in vocalization. The Journal of the Acoustical Society of America, 137(5):3005–3007, 2015. doi: 10.1121/1.4919349.

2. Hugh K Dunn. The calculation of vowel resonances, and an electrical vocal tract. The Journal of the Acoustical Society of America, 22(6):740–753, 1950.

3. Gunnar Fant. Acoustic Theory of Speech Production: With Calculations Based on X-Ray Studies of Russian Articulations. Mouton, 1971.

4. Johan Liljencrants and Gunnar Fant. Computer program for vt-resonance frequency calculations. STL-QPSR, 16:15–21, 1975.

5. Kenneth N. Stevens. Acoustic phonetics. Current studies in linguistics series, 30. MIT Press, Cambridge, Mass, 1998. ISBN 026219404X.

6. René Carré, Pierre Divenyi, and Mohamad Mrayati. Speech: A dynamic process. Walter de Gruyter GmbH & Co KG, 2017.

7. Kenneth N Stevens, Stanley Kasowski, and C Gunnar M Fant. An electrical analog of the vocal tract. The Journal of the Acoustical Society of America, 25(4):734–742, 1953.

8. Kecheng Zhang, Runhui Song, Rui Tu, Jens Edlund, Jonas Beskow, and Axel G Ekström. Modeling, synthesis and 3d printing of tube vocal tract models with a codeless graphical user interface. In Proceedings from FONETIK, pages 155–160, 2024.

9. W. L. Henke. Dynamic articulatory model of speech production using computer simulation. PhD thesis, Massachusetts Institute of Technology, 1966.

10. P. Mermelstein. Articulatory model for the study of speech production. The Journal of the Acoustical Society of America, 53(4):1070–1082, 1973.

11. Pierre Badin and Gunnar Fant. Notes on vocal tract computation. STL QPSR, 2(3):53–108, 1984.

12. Kenneth N Stevens and Arthur S House. Development of a quantitative description of vowel articulation. The Journal of the Acoustical Society of America, 27(3):484–493, 1955.

13. Uno Ingard. On the theory and design of acoustic resonators. The Journal of the Acoustical Society of America, 25(6):1037–1061, 1953.

14. J. Dang, C. H. Shadle, Y. Kawanishi, K. Honda, and H. Suzuki. An experimental study of the open end correction coefficient for side branches within an acoustic tube. The Journal of the Acoustical Society of America, 104(2):1075–1084, 1998.

15. B. Lindblom, J. Sundberg, P. Branderud, and H. Djamshidpey. On the acoustics of spread lips. Proceedings of Fonetik, 50(1):13–16, 2007.

16. Mohamad Mrayati, René Carré, and Bernard Guérin. Distinctive regions and modes: a new theory of speech production. Speech Communication, 7(3):257–286, 1988.

17. R. Song. Revisiting the third formant: Computational analysis of vocal tract constrictions using tube models and neural networks. Master’s thesis, Uppsala University, 2025.

18. Gareth (Gareth Michael) James, Daniela Witten, Trevor Hastie, and Robert Tibshirani. An introduction to statistical learning : with applications in R. Springer texts in statistics. Springer, New York, second edition edition, 2021. ISBN 9781071614174.

19. Tianqi Chen and Carlos Guestrin. Xgboost: A scalable tree boosting system. In Proceedings of the 22nd acm sigkdd international conference on knowledge discovery and data mining, pages 785–794, 2016.

20. Jerome H Friedman. Greedy function approximation: a gradient boosting machine. Annals of statistics, pages 1189–1232, 2001.

21. Diederik P Kingma and Jimmy Ba. Adam: A method for stochastic optimization. arXiv preprint 1412.6980, 2014.

22. Tadayoshi Fushiki. Estimation of prediction error by using k-fold cross-validation. Statistics and Computing, 21:137–146, 2011.

23. Sylvain Arlot and Alain Celisse. A survey of cross validation procedures for model selection. Statistics Surveys, 4, 07 2009. doi: 10.1214/09-SS054.

24. Wojciech Samek, Grégoire Montavon, Sebastian Lapuschkin, Christopher J Anders, and Klaus-Robert Müller. Explaining deep neural networks and beyond: A review of methods and applications. Proceedings of the IEEE, 109(3):247–278, 2021.

25. Scott M Lundberg and Su-In Lee. A unified approach to interpreting model predictions. In I. Guyon, U. Von Luxburg, S. Bengio, H. Wallach, R. Fergus, S. Vishwanathan, and R. Garnett, editors, Advances in Neural Information Processing Systems, volume 30. Curran Associates, Inc., 2017.

26. Lloyd S Shapley. A value for n-person games. Contributions to the Theory of Games, 2(28):307–317, 1953.

27. C. Molnar. Interpretable Machine Learning. Leanpub, 2020. ISBN 9780244768522.

28. Peter Ladefoged and Ian Maddieson. The sounds of the world’s languages. Phonological theory. Blackwell, Oxford, 1996. ISBN 0631198148.

29. Kenneth N. Stevens. On the quantal nature of speech. Journal of Phonetics, 17(1–2):3–45, 1989.

30. Hirotugu Akaike. A new look at the statistical model identification. IEEE transactions on automatic control, 19(6):716–723, 2003.

31. James Bergstra and Yoshua Bengio. Random search for hyper-parameter optimization. The journal of machine learning research, 13(1):281–305, 2012.

